# Differences in microbial community structure and soil nutrients between healthy and diseased soils of *Zingiber officinale*

**DOI:** 10.64898/2026.06.01.729252

**Authors:** Shuang Ma, Fei Fang, Jianming Li, Tie Zhang, Teng Wang

## Abstract

To investigate the differences in soil nutrients and microbial community structure in the rhizosphere between healthy and diseased *Zingiber officinale* plants, soil samples were collected from healthy root-zone soil (ZSH), healthy rhizosphere soil (RSH), diseased root-zone soil (ZSD), and diseased rhizosphere soil (RSD). Diseased soils had significantly higher pH values, whereas RSH showed the strongest acidity. Moreover, pH, AN, and AK contents in diseased soils were significantly higher than those in healthy soils, while SOM and AP were significantly lower. The α-diversity of microbial communities in diseased soils was significantly reduced, and the community structure was distinctly differentiated from that of healthy soils. In diseased soils, the abundance of potential pathogenic taxa such as *Ralstonia solanacearum* increased significantly, while beneficial genera such as *Bradyrhizobium* decreased. Redundancy analysis and correlation analysis indicated that soil pH, AN, SOM, and AP were the major environmental factors driving changes in microbial community structure. The occurrence of soil-borne diseases in *Zingiber officinale* is closely associated with soil nutrient imbalance and disruption of microbial community structure. The study identified candidate microbial taxa (e.g., beneficial *Sphingomonas, Streptomyces*) and key soil properties (pH, available nitrogen) that differentiate healthy from diseased ginger soils. Together, these findings provide a theoretical basis for improving diseased soils through microecological regulation strategies, and also serve as a foundation for generating testable hypotheses in future hypothesis-driven research on ginger soil-borne disease suppression.

## Introduction

*Zingiber officinale* is an economically important crop in China, widely used for culinary, medicinal, and cosmetic purposes due to its high gingerol content, rich flavor, and medicinal properties (Goyal et al. 2021). Shandong, Yunnan, and Hunan provinces are known for their diverse local ginger varieties. However, ginger production is severely threatened by bacterial wilt, a disease caused by *Ralstonia solanacearum* (Li et al. 2021, Zhang et al. 2022). In recent years, the continuous expansion of ginger cultivation and the widespread practice of continuous cropping have exacerbated soil-borne diseases. Frequent outbreaks of bacterial wilt have led to substantial yield losses and quality deterioration, as well as pesticide overuse and soil degradation, thereby constraining the sustainable development of the ginger industry (Liu et al. 2023).

Soil-borne diseases fundamentally arise from rhizosphere dysbiosis, characterized by nutrient imbalance and disruption of microbial community structure (Wei et al. 2016). In ginger cultivation, such disorders have been documented to cause yield losses of 40-50% in epidemic regions, with localized cases of complete crop failure (Feng et al. 2022). A representative example is the bacterial wilt (*Ralstonia solanacearum*) outbreak in Yanshan County, a major ginger-producing region in southeastern Yunnan Province. These findings underscore the necessity to elucidate environment-microbiome interactions in diseased versus healthy soils.

Rhizosphere microorganisms play a key role in nutrient cycling and disease suppression, and their diversity and abundance are important indicators of soil health (Bakker et al. 2018, Fan et al. 2021). In healthy soils, complex and stable microbial communities occupy ecological niches, suppressing pathogens through competition, antagonism, and induced resistance (Kumar et al. 2022, Kwak et al. 2022). Conversely, continuous cropping or improper fertilization can alter soil physicochemical properties, reduce microbial diversity, and shift community structure toward pathogen enrichment, thereby weakening soil suppressiveness and promoting disease outbreaks (Deng et al. 2023). For example, *Fusarium oxysporum* abundance increases while beneficial *Pseudomonas* decreases in *Fusarium* wilt-diseased soils (Li et al. 2023). However, the linkage between ginger rhizosphere microbiota and plant health remains poorly understood, warranting systematic comparison of soil properties and microbial communities between healthy and diseased soils.

Soil nutrients fundamentally shape rhizosphere microbiome assembly through trophic and signaling pathways. Beyond their roles as plant macronutrients, nitrogen, phosphorus, and potassium dynamically reconfigure microbial consortia (Zhang et al. 2021, Liu et al. 2022, Zhou et al. 2023). In the soil microenvironment, pH is a primary driver of bacterial community structure, as it affects cell membrane charge, enzyme activity, and nutrient availability (Lauber et al. 2009, Bahram et al. 2018). Soil organic matter and available nitrogen directly determine microbial resource availability, with organic-rich soils supporting more heterotrophic microorganisms (Liang et al. 2017, Chen et al. 2021). Therefore, these environmental factors collectively determine the composition of the rhizosphere microbiome.

Research on ginger soil-borne diseases has largely overlooked the rhizosphere microbiome, focusing instead on pathogen isolation and chemical control. We collected rhizosphere and root-zone soils from healthy and diseased ginger plants, measured physicochemical parameters (pH, AN, SOM, AK, AP), and performed high-throughput sequencing to compare bacterial community structure. Using redundancy analysis and Spearman correlation, we aimed to identify key environmental factors driving microbial shifts and elucidate the microecological mechanisms of ginger bacterial wilt. Our findings will support green disease control, soil microecological remediation, evidence-based fertilization, and the sustainable development of the ginger industry.

## Materials and methods

### Soil sample preparation

Soil samples were collected from a ginger wilt disease-affected field in Zhela Township, Yanshan County, Wenshan Prefecture, Yunnan Province, China (23°40′03″N, 104°28′29″E). A total of 12 soil samples were collected, including healthy rhizosphere soil (RSH), diseased rhizosphere soil (RSD), healthy root-zone soil (ZSH), and diseased root-zone soil (ZSD), each taken from the corresponding plant’s root zone or rhizosphere. The collected soils were immediately placed into sterile ziplock bags, transported to the laboratory in an icebox at low temperature. Each fresh soil sample was divided into two portions. One portion was used for microbial analysis: fresh soil was passed through a 2 mm sterile sieve (the sieve was disinfected with 75% alcohol and air-dried), and after removing plant residues and stones, the soil was stored at -80°C for further use. The other portion was used for routine chemical analysis: soil was air-dried, ground, and sieved through 2 mm mesh sieves to determine soil pH, alkaline hydrolysis nitrogen (AN), available phosphorus (AP), available potassium (AK), and soil organic matter (SOM).

### Determination of soil alkaline hydrolyzable nitrogen

Soil hydrolyzable nitrogen was determined using the alkaline hydrolysis diffusion method (Lu et al. 2023). 2.00 g of air-dried soil was placed in the outer chamber of a Conway cell, and after adding the reducing agent (for upland soil) and 10 mL of NaOH, the cell was sealed and incubated at 40°C for 24 h. The liberated NH_3_ was trapped in 2 mL of 20 g·L^−1^ boric acid-indicator solution and titrated with 0.01 mol·L^−1^ HCl. Results are expressed in mg·kg^−1^, with parallel determination error ≤10%.

### Determination of soil available phosphorus

Soil available phosphorus was extracted using an ammonium chloride-hydrochloric acid extractant at a soil-to-solution ratio of 1:10 (Bray and Kurtz, 1945). The mixture was shaken at 22°C for 30 min, then filtered. An aliquot of the filtrate was mixed with boric acid solution to adjust the acidity, followed by color development using the molybdenum-antimony-ascorbic acid chromogenic reagent.

### Determination of soil available potassium

An air-dried soil sample (5.00 g) was mixed with 50 mL of 1 mol·L^−1^ ammonium acetate solution (Hanway and Heidel, 1952). The mixture was shaken for 30 min and then filtered. The filtrate was directly used to determine potassium content using a flame photometer.

### Determination of soil pH

An air-dried soil sample (10.0 g) passing through a 2 mm sieve was mixed with 25 mL of CO_2_-free water (soil-to-water ratio of 1:2.5). The suspension was stirred for 1 min, allowed to stand for 30 min, and then the pH was measured using a pH meter (precision of 0.01).

### Determination of soil organic matter

Soil organic matter was determined by the K_2_Cr_2_O_7_-oil bath heating method. An air-dried soil sample (0.5 g) was placed into a hard glass tube, mixed with 10.00 mL of 0.4 mol·L^−1^ K_2_Cr_2_O_7_-H_2_SO_4_ solution, and digested in an oil bath at 170–180°C for 5 min. After cooling, the digest was transferred and the residual K_2_Cr_2_O_7_ was titrated with FeSO_4_·(NH_4_)_2_SO_4_ standard solution using ferron as indicator. Two blanks (ignited pumice powder instead of soil) were included per batch.

### Sample DNA extraction, library construction, and sequencing

Total genomic DNA of the microbial community in soil samples was extracted following the manufacturer’s instructions of the E.Z.N.A.® soil DNA kit (Omega Bio-tek, Norcross, GA, USA). After extraction, DNA concentration and purity were measured, and DNA integrity was checked by 1% agarose gel electrophoresis. The DNA was fragmented using a Covaris M220, and fragments of approximately 350 bp were selected for paired-end library construction. Library preparation was performed using the NEXTFLEX Rapid DNA-Seq library preparation kit (Bioo Scientific, USA). Metagenomic sequencing was conducted on the Illumina NovaSeq™ X Plus platform (Shanghai Majorbio Bio-pharm Technology Co., Ltd.).

### Quality control and assembly

A total of 12 samples were collected, yielding 137.61 Gb of raw data. Raw reads were filtered using fastp (version 0.20.0) to remove low-quality bases, and then host sequences were aligned and removed using BWA (version 0.7.17). After quality control, 125.87 Gb of high-quality clean data were obtained, with an average of 10.49 Gb per sample. Clean reads from each sample were assembled using MEGAHIT (version 1.2.9), and contigs shorter than 500 bp were discarded. Open reading frames (ORFs) were predicted from the assembled sequences using Prodigal (version 2.6.3), and sequences with a length ≥ 100 bp were retained.

### Non-redundant gene set construction and abundance calculation

The predicted gene sequences from all samples were clustered and de-redundantly assembled using CD-HIT (version 4.6.1) with parameters set to identity ≥ 90% and coverage ≥ 90%, to construct a non-redundant gene set. High-quality reads from each sample were then aligned to the non-redundant gene set using SOAPaligner (version soap2.21 release) with identity ≥ 95%, and the abundance information of genes in the corresponding samples was calculated.

### Statistical analysis

Alpha diversity and beta diversity analyses were performed using R language (version 4.2.0). Principal coordinate analysis (PCoA) based on Bray-Curtis distance, combined with PERMANOVA (Adonis, 999 permutations), was used to assess overall differences in microbial community structure among samples. Wilcoxon rank-sum test (for two-group comparisons) or Kruskal-Wallis test (for multi-group comparisons) was applied for differential analysis between groups. LEfSe analysis (linear discriminant analysis effect size) with an LDA threshold > 2 was used to screen differential species. Spearman correlation analysis was conducted to evaluate associations between microbial species and physicochemical factors, and random forest algorithm was employed to identify biomarkers that distinguish different groups. Redundancy analysis (RDA) was used to assess the relationship between microbial communities and physicochemical factors, and variance partitioning analysis (VPA) was performed using the vegan package in R language to quantify the contribution of each environmental factor to community variation.

## Results

### Differences in physicochemical properties between healthy and diseased soil

There was a significant differentiation in pH values between healthy and diseased ginger soils (P < 0.05). The soil pH followed the order: RSD > ZSD > ZSH > RSH. Among them, the RSD treatment had the highest pH, with an average value of 5.3675, significantly higher than all other treatments; the ZSD treatment ranked second (5.2450), which was higher than ZSH and RSH; the ZSH treatment also had a higher pH than RSH (P < 0.05). The average pH of healthy soils (ZSH, RSH) was 4.81, which is in the slightly acidic range. In contrast, the average pH of diseased soils (ZSD, RSD) increased to 5.31, significantly higher than that of healthy soils (P < 0.05). This trend indicates that the disease led to an increasing trend in rhizosphere soil pH.

The alkaline hydrolyzable nitrogen (AN) content in diseased soils showed a strong accumulation effect. Soil AN content followed the order: ZSD > RSD > ZSH > RSH. The average AN concentrations in diseased soils ZSD and RSD reached 289.33 mg/kg and 282.33 mg/kg, respectively, more than double those in healthy soils (ZSH: 128.33 mg/kg; RSH: 93.33 mg/kg). This reflects that excessive nitrogen accumulation in diseased soils favors disease development.

Soil available potassium (AK) showed a trend consistent with that of alkaline hydrolyzable nitrogen. The average AK concentration in diseased soils was also significantly higher than that in healthy soils, with the highest value (231.57 mg/kg) observed in the ZSD treatment. The overall order was ZSD > ZSH > RSD > RSH. Although potassium is an essential element for plant growth, a high-potassium environment may induce succession in the soil microbial community structure. Moreover, excessively high potassium ions can antagonize the uptake of cations such as calcium and magnesium, causing physiological metabolic disorders in plants and potentially indirectly exacerbating disease incidence.

Soil available phosphorus (AP) showed the opposite trend to nitrogen and potassium, following the order: ZSH > RSH > ZSD > RSD. The average AP concentration in healthy soils was significantly higher than that in diseased soils. Among them, the healthy soil ZSH had the highest AP concentration (19.74 mg/kg), which was closest to 20 mg/kg, whereas AP in diseased soils was below 5 mg/kg, indicating severe phosphorus deficiency. The reason is that phosphorus is a core element in the composition of ATP, nucleic acids, and other life substances. Phosphorus deficiency not only limits the growth and development of ginger plants but also hinders the activation of phosphorus by soil microorganisms, thereby impeding the material cycling in the rhizosphere microecosystem.

Soil organic matter (SOM) concentration in healthy soils was stable at approximately 35 mg/kg, while in diseased soils ZSD and RSD it decreased significantly to 28.16 mg/kg and 23.87 mg/kg, a reduction of about 26%. Studies indicate that organic matter deficiency implies a severe shortage of carbon sources for soil microorganisms, which may lead to reduced metabolic activity of the soil microbial community. Meanwhile, the decline in organic matter content weakens the formation of soil aggregate structure, reduces soil water and nutrient retention capacity, and diminishes the ability of ginger plants to survive under environmental stress (Figure 1).

**Figure 1.**
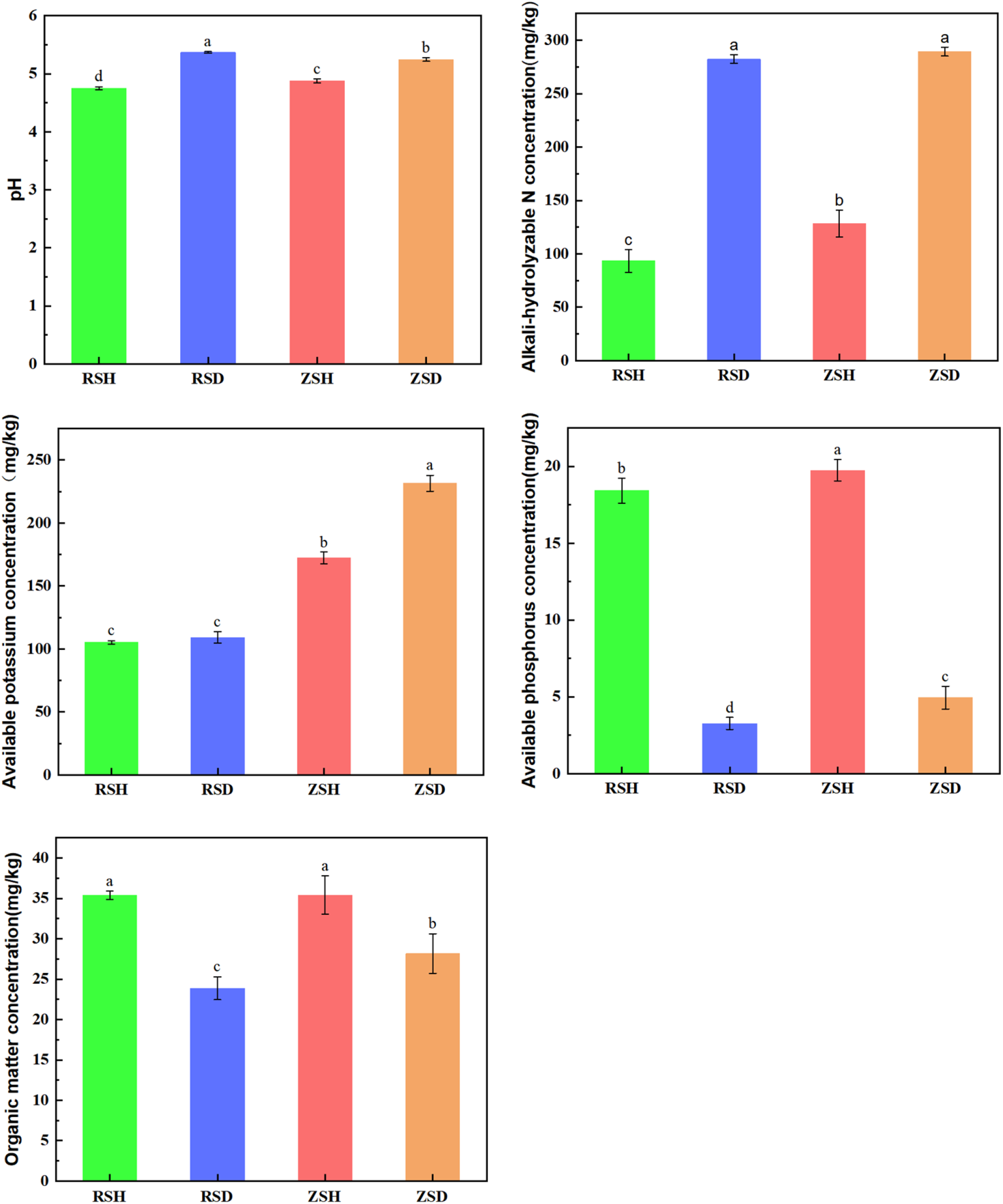
Physicochemical properties of healthy and diseased soil.

In summary, diseased soils exhibited a nutrient imbalance characterized by elevated pH, excessive nitrogen and potassium, and decreased organic matter and available phosphorus, which created a physicochemical environment conducive to the occurrence of soil-borne diseases.

Alpha diversity analysis showed that the Ace index of healthy soil groups (ZSH, RSH) was significantly higher than that of diseased soil groups (ZSD, RSD) (P = 0.02353). Specifically, there was a significant difference in alpha diversity among the different groups, with the Ace index of the RSH and ZSH groups being significantly higher than that of the RSD and ZSD groups (Figure 2-A). This indicates that the rhizosphere microecosystems of the RSH and ZSH groups harbor higher soil bacterial species richness and better community diversity.

**Figure 2.**
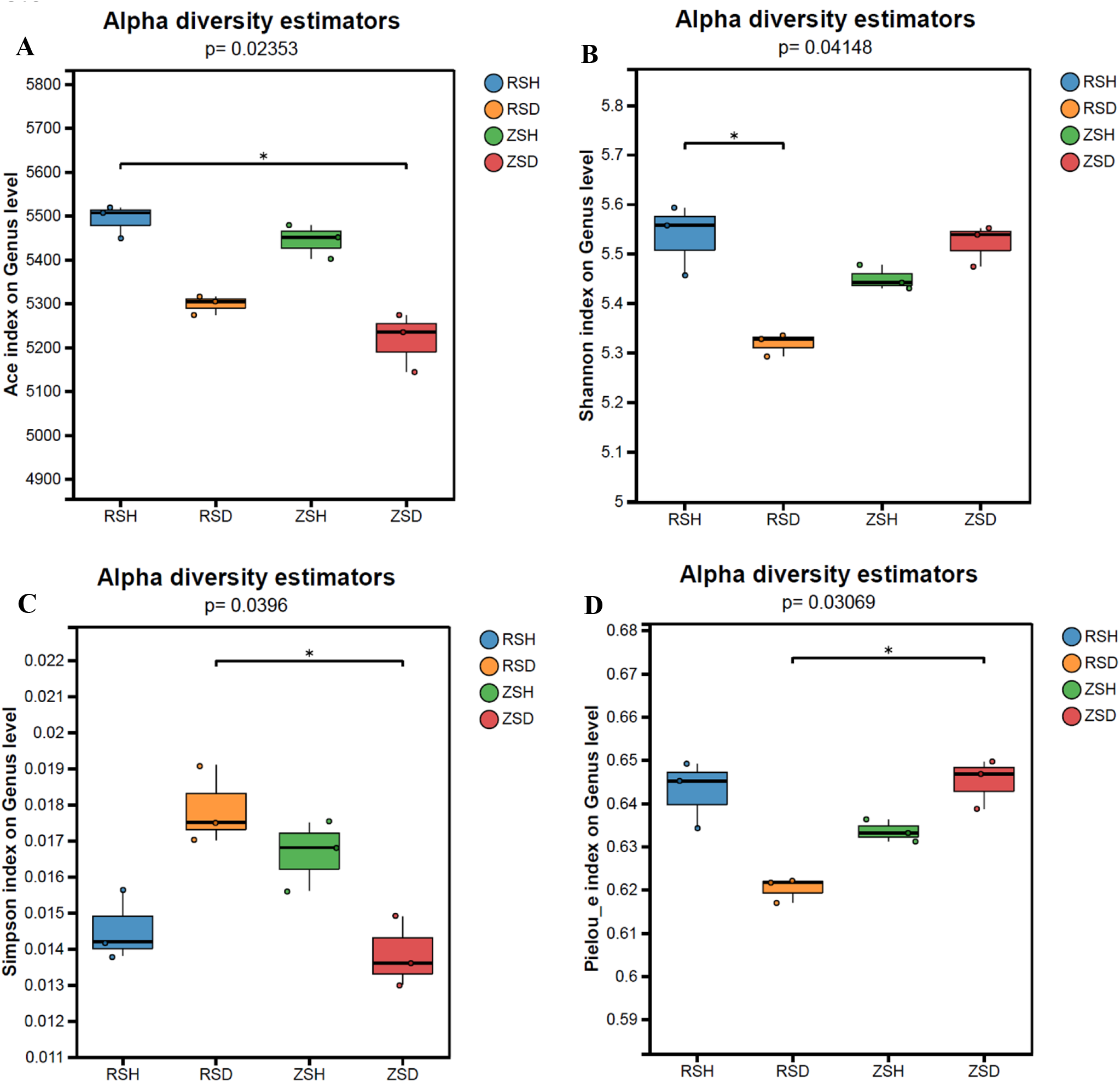
Alpha diversity indices of bacterial communities in different soil groups. A: Ace index; B: Shannon index; C: Simpson index; D: Pielou evenness index (Pielou-e).

Shannon index analysis showed a significant difference between healthy rhizosphere soil (RSH) and diseased rhizosphere soil (RSD) (p=0.04148), with the shannon index of the RSH group being significantly higher than that of the RSD group. In contrast, no significant difference was observed between healthy root-zone soil (ZSH) and diseased root-zone soil (ZSD) (p>0.05). Additionally, no significant differences were detected between RSH and ZSH, nor between RSD and ZSD (p>0.05) (Figure 2-B). These results indicate that the bacterial community diversity in rhizosphere soil is more sensitive to the occurrence of *Z. officinale* disease.

Simpson index analysis showed a significant difference between diseased rhizosphere soil (RSD) and diseased root-zone soil (ZSD) (p=0.0396), with the simpson index of the RSD group being significantly higher than that of the ZSD group. No significant differences were detected among the remaining treatment pairs (p > 0.05) (Figure 2-C). This result indicates that under disease conditions, the dominance structure of bacterial communities in rhizosphere and root-zone soils diverged significantly.

Pielou evenness index (Pielou-e) analysis showed a significant difference between diseased rhizosphere soil (RSD) and diseased root-zone soil (ZSD) (p=0.03069), with the Pielou-e index of the RSD group being significantly lower than that of the ZSD group. No significant differences were detected between RSD or ZSD and healthy rhizosphere soil (RSH) or healthy root-zone soil (ZSH) (p>0.05) (Figure 2-D). This finding suggests that under disease pressure, bacterial community evenness in the rhizosphere and root-zone soils diverges markedly.

Healthy soil samples (ZSH and RSH) were closely clustered together with strong intra-group reproducibility, indicating that the rhizosphere bacterial community structure of healthy plants exhibited high consistency (Figure 3). In contrast, diseased soil samples (ZSD and RSD) gathered on the opposite side and were significantly separated from the healthy soil group along the PC1 axis (R=0.849, P=0.001). These results suggest that plant health status is a crucial factor driving differences in the rhizosphere microbial community structure of *Z. officinale*. The occurrence of disease not only alters the abundance of individual microbial species but also reshapes the overall composition of the rhizosphere microbial community.

**Figure 3.**
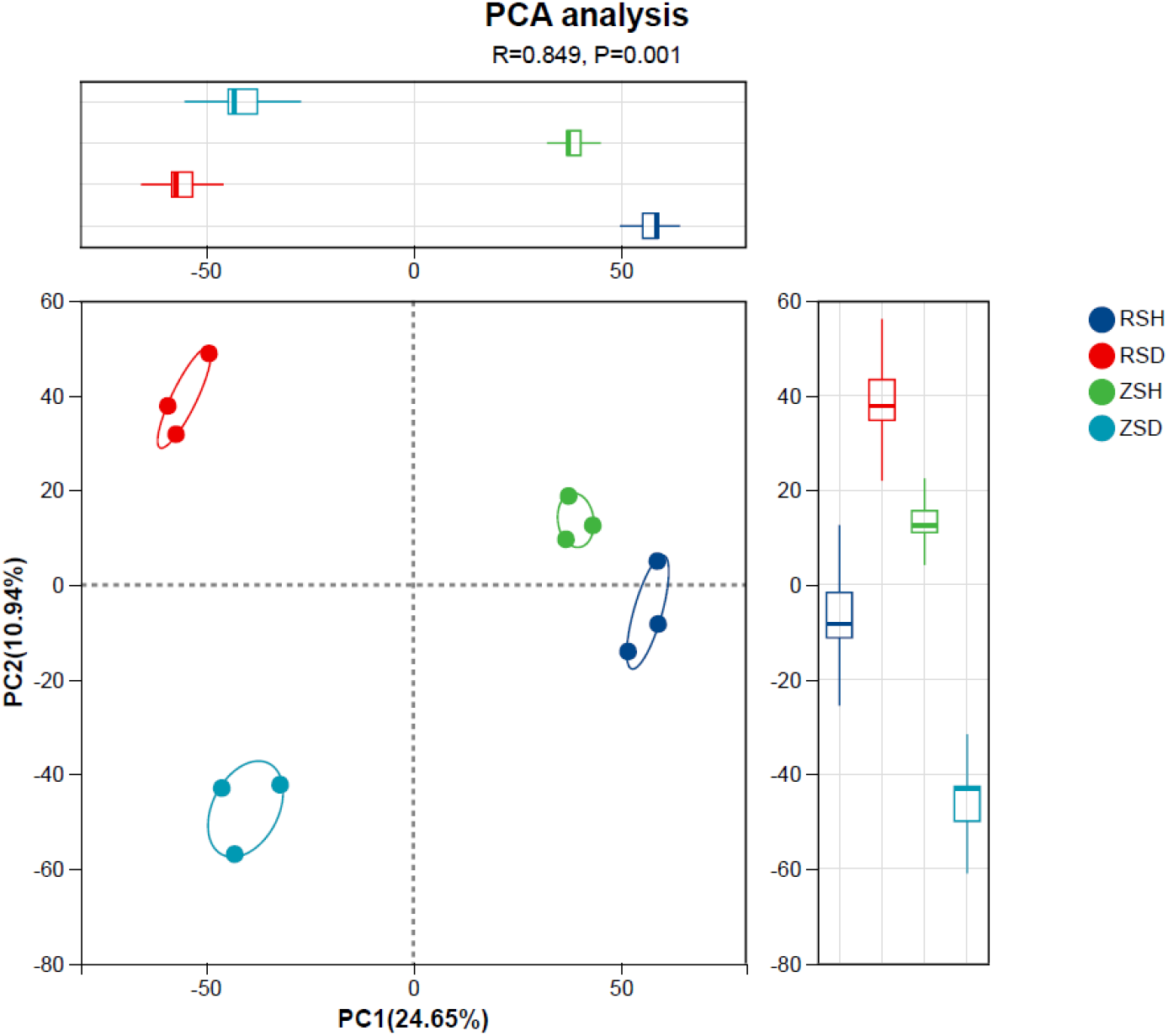
β-diversity of soil samples.

At the genus level (Figure 4), the composition of dominant bacterial genera in healthy and diseased soils was similar, with *Sphingomonas, Bradyrhizobium*, and *Ralstonia* ranking as the top three. However, their relative abundances showed significant differences. Among them, the abundance of the pathogenic genus *Ralstonia* (the bacterial wilt pathogen) was markedly increased in the rhizosphere soil of diseased plants, forming a substantial difference compared with healthy soil samples. Notably, a high enrichment of *Ralstonia* was observed in the diseased soil sample RSD.

**Figure 4.**
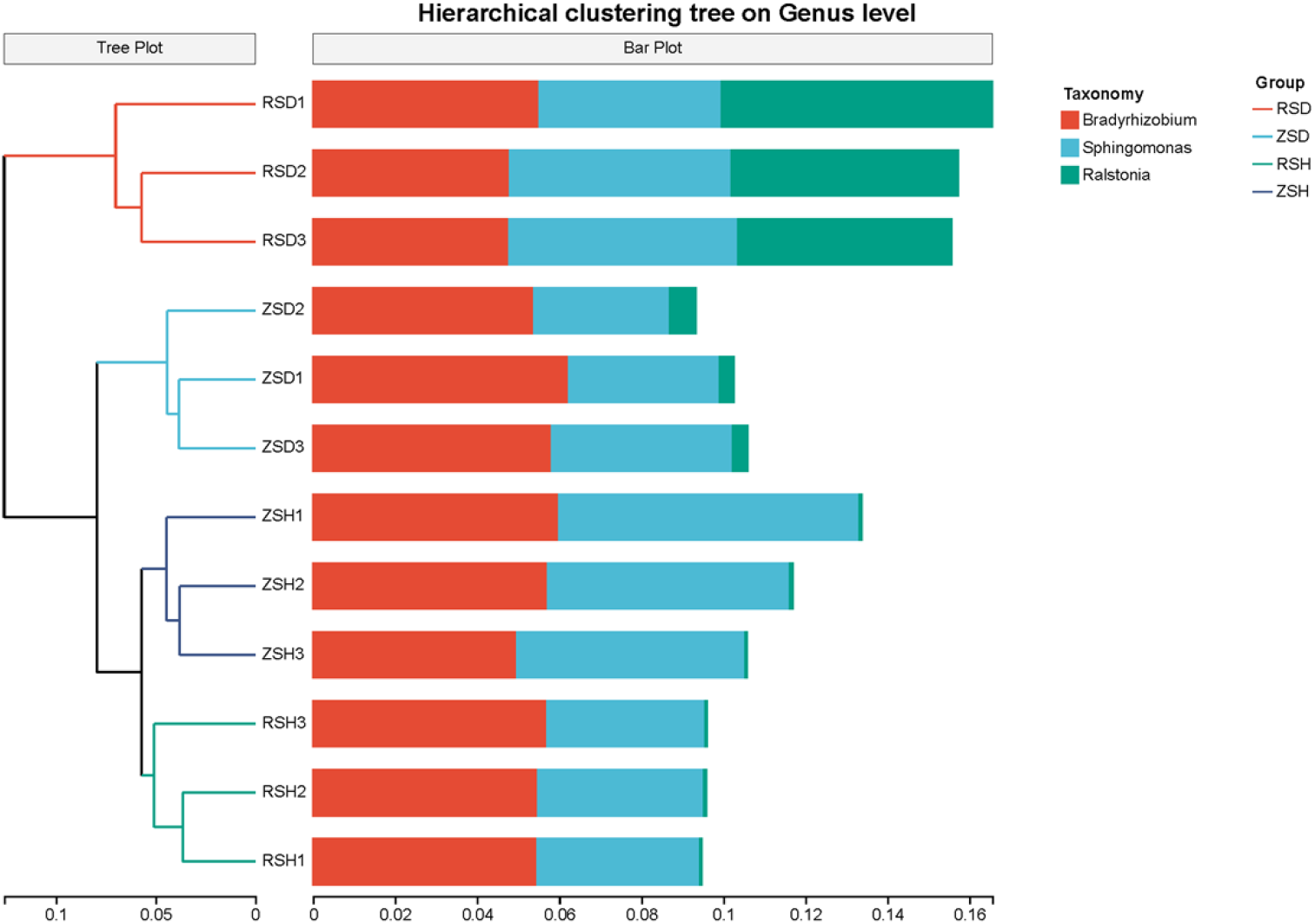
Relative abundance (top 3) and hierarchical clustering analysis of microbial communities at the genus level in different soil samples.

The species abundance heatmap (Figure 5) further quantified the distribution patterns of dominant microbial communities, revealing significant differences in bacterial abundance among groups. Notably, the abundance of *Ralstonia* (the bacterial wilt pathogen) in the RSD group was higher than that in other groups. Obvious color gradient changes can be observed in the heatmap, in the modules of diseased soils (RSD and ZSD), taxa such as *Ralstonia* and *Bradyrhizobium* appeared in red, indicating higher enrichment, suggesting that these microbial groups may be indicator species for disease occurrence. In contrast, the modules specific to healthy soils (RSH and ZSH) exhibited a distinct characteristic enrichment pattern, with taxa such as *Sphingomonas* and *Ktedonobacter* showing red coloration and higher enrichment. This strikingly different heatmap pattern establishes a microbial “fingerprint” that distinguishes healthy from diseased soils, indicating that the health status of *Z. officinale* is highly correlated with the presence or absence of specific functional microbial groups in the rhizosphere.

**Figure 5.**
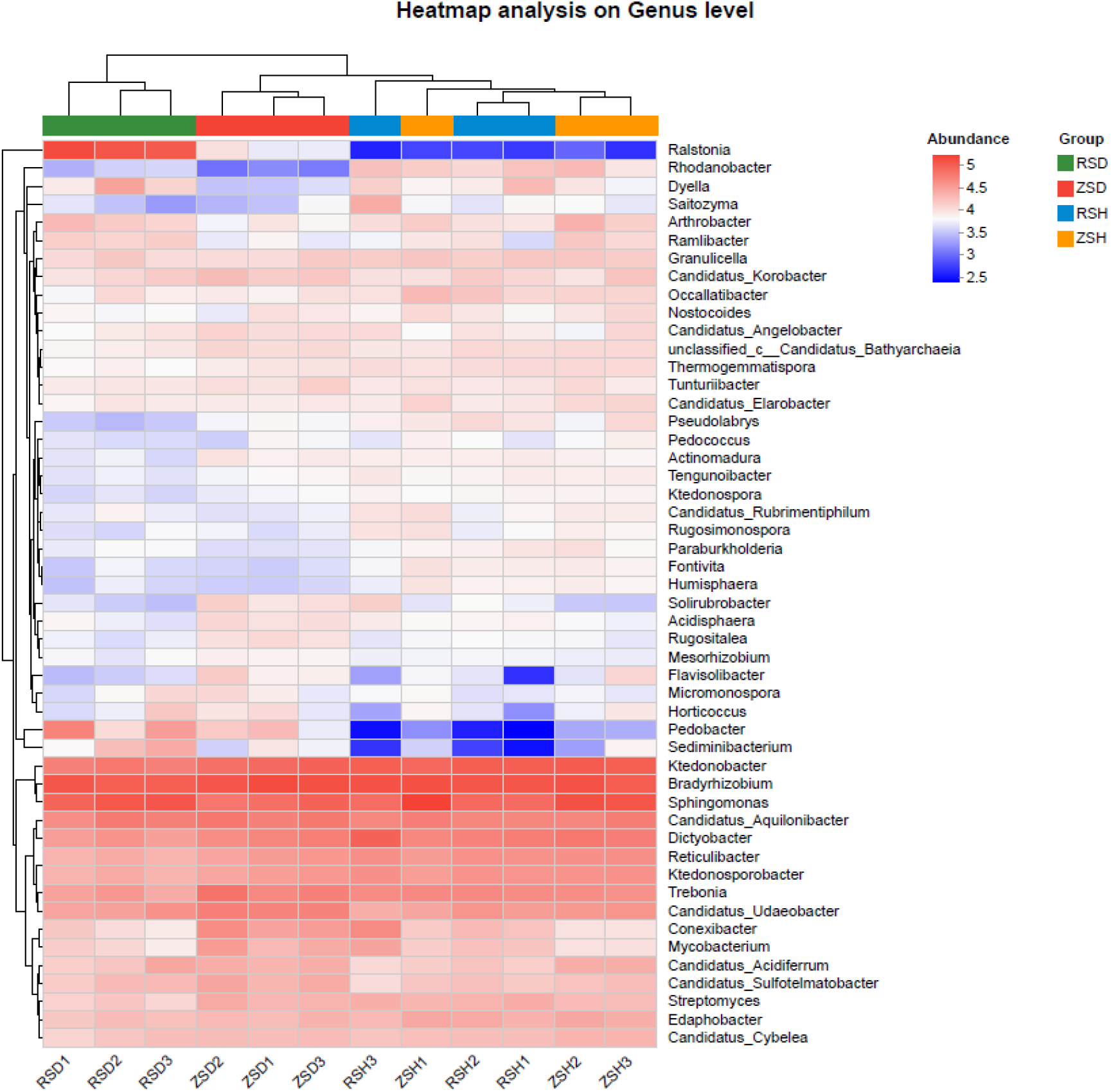
Relative abundance heatmap of dominant genera in different soil samples.

Differential genus abundance analysis (Figure 6) revealed that the relative abundances of the ten selected key genera differed significantly among the four treatment groups, rhizosphere soil of healthy plants (RSH), rhizosphere soil of diseased plants (RSD), root-zone soil of healthy plants (ZSH), and root-zone soil of diseased plants (ZSD) (P<0.05). Among these, *Ralstonia* showed the most significant intergroup difference (P=0.02374), while *Dictyobacter* exhibited a relatively lower significance level (P=0.04344). Overall, the abundance patterns of the microbial community were significantly co-regulated by plant health status and soil microhabitat.

**Figure 6.**
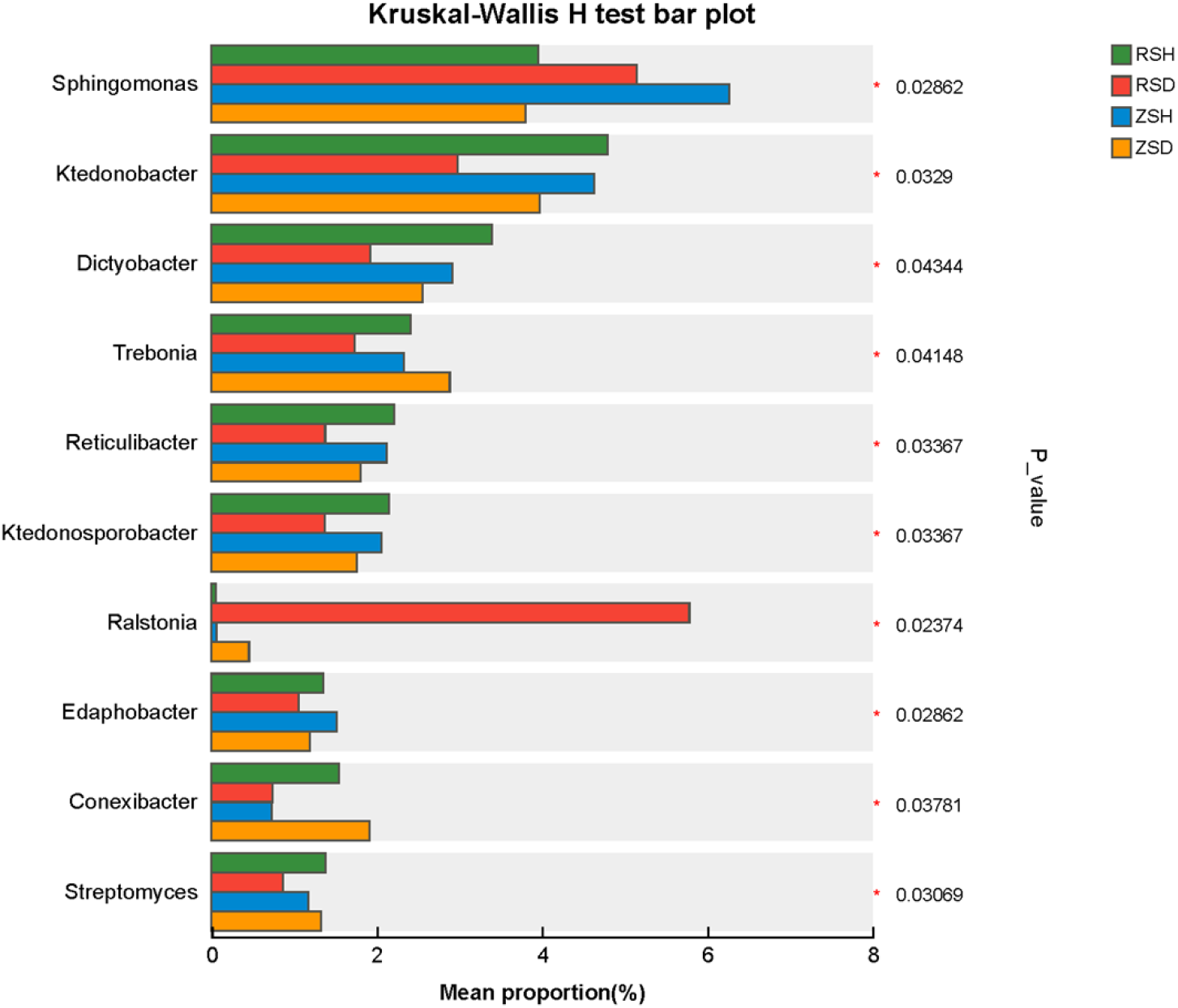
Genus-level differential species abundance plot based on the Kruskal–Wallis test.

In terms of disease-associated microbiota, *Ralstonia* displayed a strong disease-specific rhizosphere enrichment. Its relative abundance was highest in the rhizosphere soil of diseased plants (RSD), significantly higher than that in healthy rhizosphere, healthy root-zone, and diseased root-zone soils, while it was present at very low abundance in non-diseased rhizosphere environments. This indicates that *Ralstonia* primarily colonizes massively the rhizosphere microhabitat of diseased plants and serves as a key indicator genus for soilborne disease incidence. Moreover, its enrichment exhibits strict rhizosphere specificity, with no significant accumulation of the pathogen observed in root-zone soil.

Regarding beneficial microbiota associated with healthy roots, genera such as *Sphingomonas, Streptomyces, Ktedonobacter*, and *Dictyobacter* were significantly enriched in the rhizosphere soil of healthy plants (RSH), and their abundances were markedly reduced in diseased rhizosphere soil. Among them, *Sphingomonas* was the most dominant genus across all samples, maintaining high abundance in healthy rhizosphere and healthy root-zone soils but showing a clear decline in the diseased rhizosphere environment. Likewise, *Streptomyces*, a typical biocontrol bacterium, followed the same distribution pattern-enriched in the healthy rhizosphere and depleted in the diseased rhizosphere-indicating that the rhizosphere of healthy plants can selectively recruit beneficial functional microbiota, thereby constructing a rhizosphere microecological environment with disease-suppression potential.

For low-abundance differential genera, taxa such as *Trebonia, Reticulibacter, Ktedonosporobacter, Edaphobacter*, and *Conexibacter* exhibited generally low relative abundances but still showed significant intergroup differences. These genera were mostly distributed specifically in either diseased root-zone soil or healthy rhizosphere soil, suggesting that disease occurrence not only alters the structure of core rhizosphere microbial communities but also significantly affects the microbial community composition of non-rhizosphere (root-zone) soil, leading to an imbalance in soil microecological flora structure.

Redundancy analysis (RDA) revealed the driving mechanisms of environmental factors on bacterial community structure variation. The RDA results (Figure 7) showed that the first two axes collectively explained 78.64% of the total variation in the bacterial community, with RDA1 explaining 65.19% and RDA2 explaining 13.45%, indicating that the selected environmental factors were strongly representative. Soil pH, alkali-hydrolyzable nitrogen (AN), soil organic matter (SOM), available phosphorus (AP), and available potassium (AK) were the main environmental factors driving differences in bacterial community composition. The vector directions of soil pH, AN, and AK pointed toward the diseased soil samples (ZSD and RSD), indicating positive correlations with diseased soils. Among these, pH had the longest vector length and exerted the greatest influence, suggesting that higher levels of these factors progressively shifted the soil microenvironment toward a diseased state. In contrast, the vector directions of SOM and AP pointed toward the healthy soil samples (ZSH and RSH), indicating positive correlations with healthy soils. Among these, SOM had the greatest influence, suggesting that these factors are key positive drivers for maintaining a healthy soil microecology. Taken together, these results clearly demonstrate that soil nutrient variation is the core driving force behind the succession of microbial community structure.

**Figure 7.**
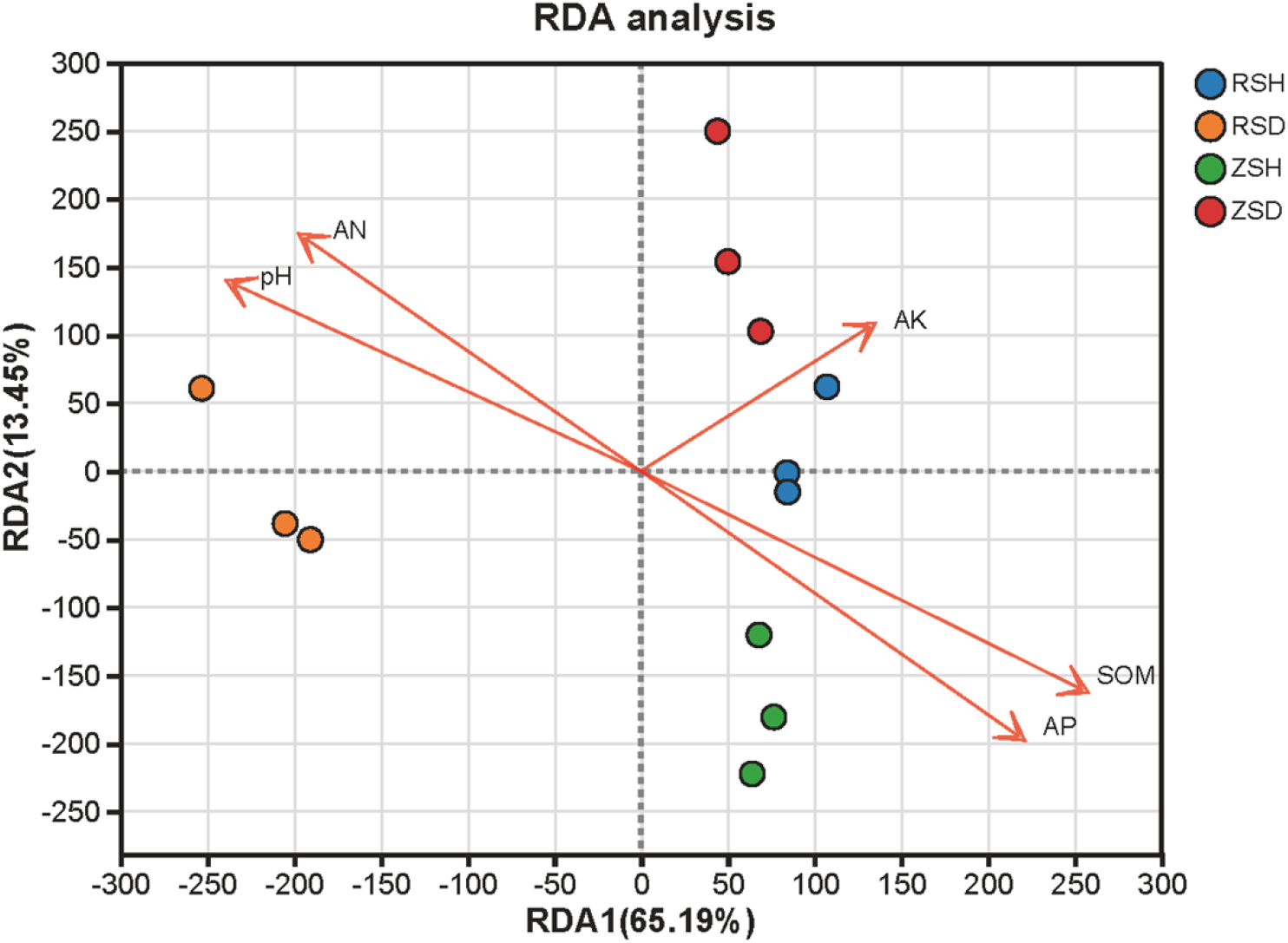
Redundancy analysis (RDA) of soil microbial community structure and environmental factors.

Spearman correlation heatmap analysis quantified the strength of relationships between nutrient factors and specific microbial groups (Figure 8), further validating the associations between environmental factors and bacterial genera, and confirming that environmental factors drive differences in community structure by regulating genus abundances. SOM showed significant positive correlations with *Bradyrhizobium* and *Sphingomonas* in healthy soils, while available phosphorus (AP) exhibited significant positive correlations with *Flavisolibacter* and *Turnturiibacter* in healthy soils (P < 0.05). This suggests that increasing soil organic matter and available phosphorus supply is an effective approach for promoting beneficial microbial communities. Soil pH and alkali-hydrolyzable nitrogen (AN) showed significant positive correlations with the potential pathogens *Ralstonia*. This indicates that diseased soils are typically characterized by a high-nitrogen, slightly alkaline environment, which may provide optimal nutritional and physical conditions for pathogen proliferation. Together, these results demonstrate that soil nutrients shape the rhizosphere bacterial community structure through direct selective effects and indirect metabolic regulation, ultimately influencing the health status of ginger plants.

**Figure 8.**
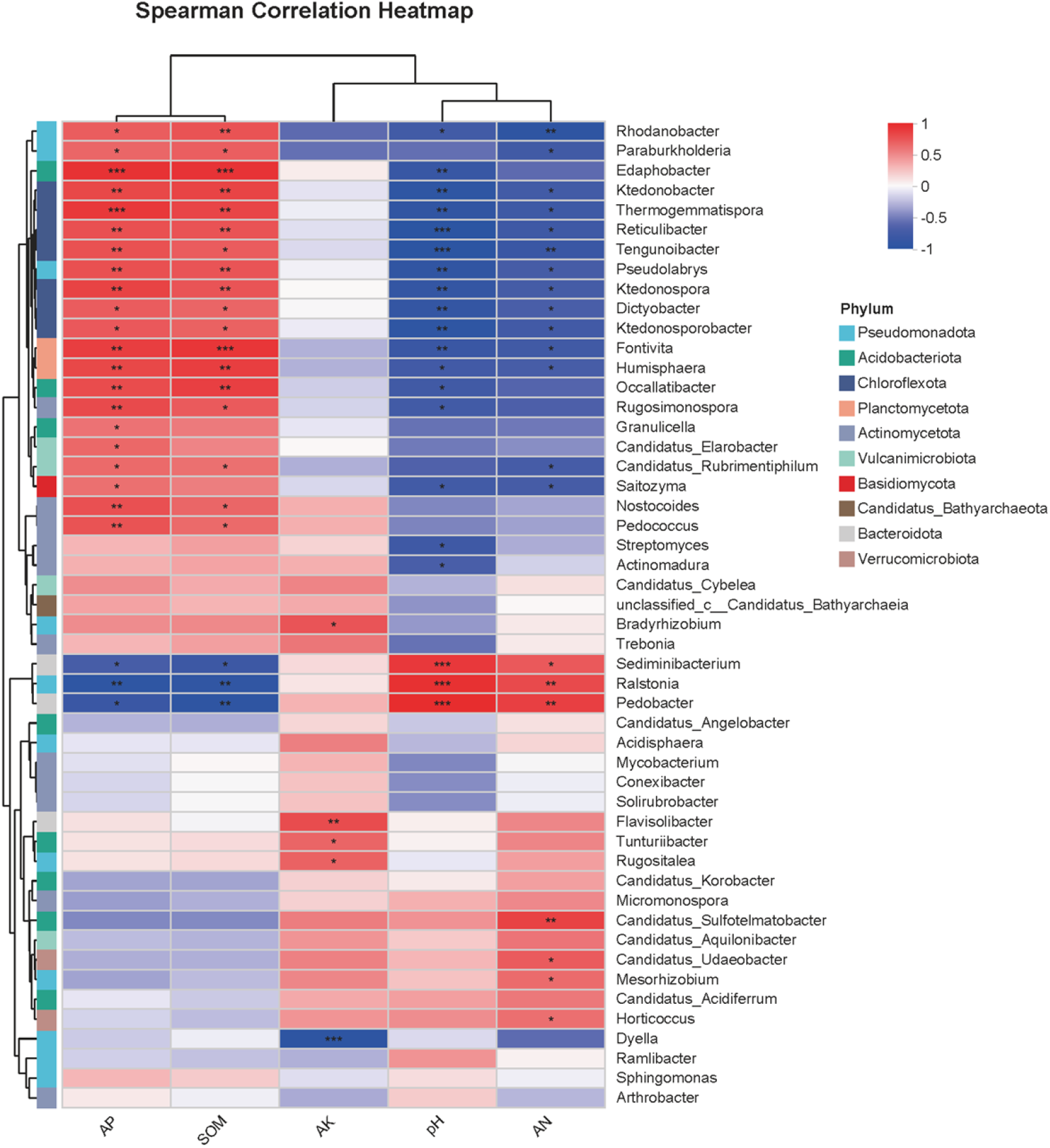
Spearman correlation heatmap of soil microbial genera and environmental factors. Note: The x-axis represents environmental factors (AP: available phosphorus, SOM: soil organic matter, AK: available potassium, pH: pH value, AN: alkali-hydrolyzable nitrogen), and the y-axis represents microbial genera. The legend on the left indicates the phylum-level classification of each genus using different colors. The color of each cell represents the Spearman correlation coefficient (red: positive correlation, blue: negative correlation), with darker colors indicating stronger correlations. * indicates significant correlation (p < 0.05), ** indicates very significant correlation (p < 0.01), *** indicates highly significant correlation (p < 0.001).

In this study, network analysis was performed to reveal the association patterns between the target bacterial community and soil environmental factors (Figure 9). The results showed that significant correlations existed between the 7 bacterial genera and 5 soil physicochemical factors. Among them, pH, alkali-hydrolyzable nitrogen (AN), available phosphorus (AP), and soil organic matter (SOM) were key drivers shaping the community structure, while available potassium (AK) was only significantly and positively correlated with *Bradyrhizobium*. Specifically, *Ralstonia* was significantly positively correlated with pH and AN, and significantly negatively correlated with AP, acting as a key hub connecting multiple environmental factors. In addition, significant synergistic relationships were observed among some bacterial genera; for example, *Ralstonia* was significantly positively correlated with *Ktedonosporobacter*, indicating a complex interaction network within the community.

**Figure 9.**
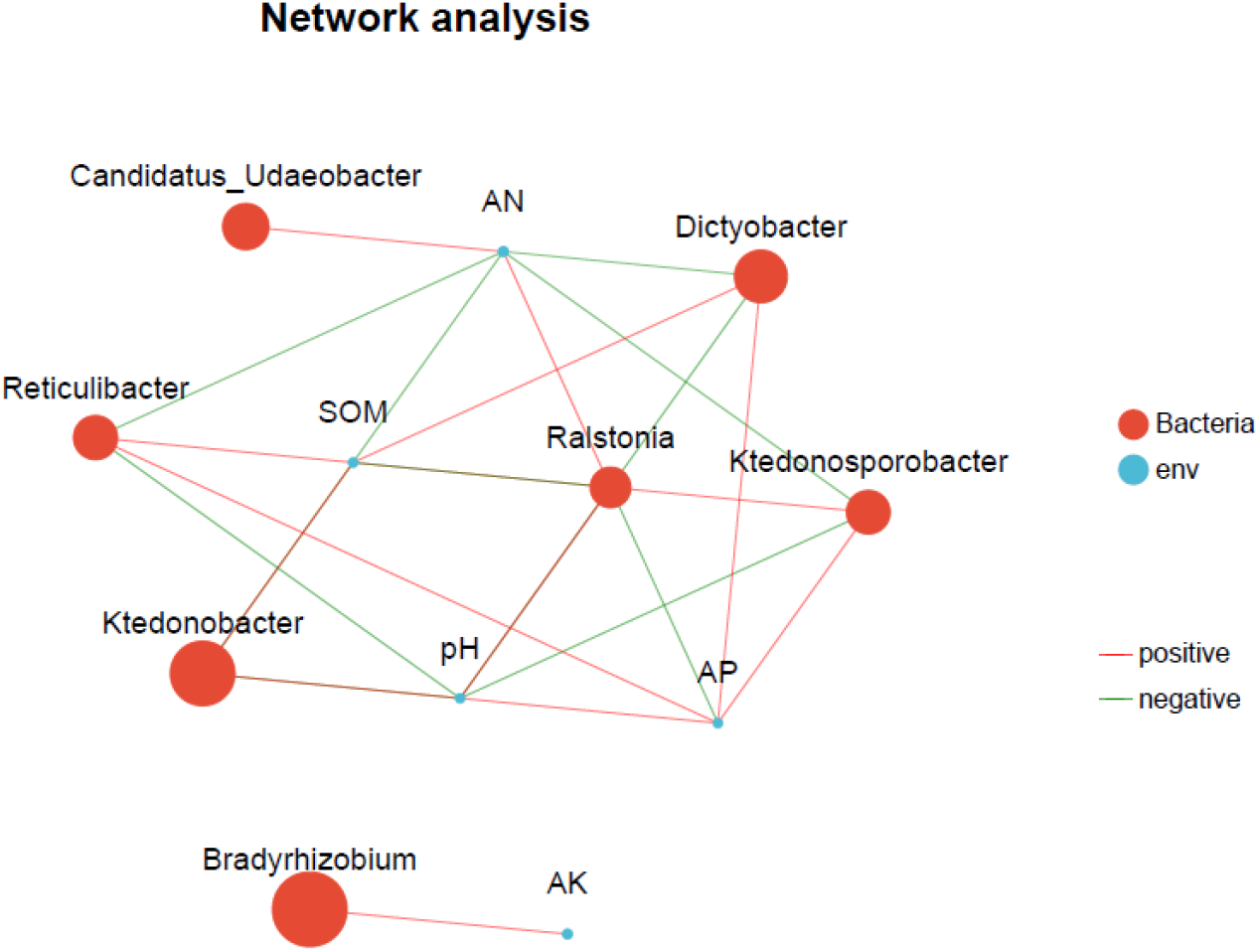
Association network analysis of target bacterial communities and soil environmental factors.

## Discussion

The results of this study showed that soils from *Z. officinale* bacterial wilt-diseased plots and healthy plots exhibited significant differences in both physicochemical properties and microbial community structure, indicating that the occurrence of soilborne disease is closely associated with soil nutrient imbalance and microbial community dysbiosis. The rhizosphere soil of diseased ginger plants displayed a characteristic nutrient imbalance, including elevated pH, excessive accumulation of alkali-hydrolyzable nitrogen and available potassium, and markedly lower levels of soil organic matter and available phosphorus. Excessive or unbalanced application of nitrogen and potassium fertilizers is likely the primary cause of this nutrient imbalance. High nitrogen input not only directly increases soil alkali-hydrolyzable nitrogen content but also indirectly raises soil pH by altering root exudate composition and nitrification processes (Gu et al. 2021). Meanwhile, excessive potassium accumulation further disrupts soil cation balance, thereby affecting the microbial living environment (Saba et al. 2024). In addition, continuous depletion of soil organic matter and the easy fixation or loss of available phosphorus lead to carbon source scarcity and insufficient phosphorus supply, significantly reducing soil buffering capacity as well as nutrient retention and supply capabilities. This systemic nutrient imbalance fundamentally alters the rhizosphere micro-ecological environment, creating favorable conditions for pathogen proliferation while simultaneously inhibiting the growth of beneficial microorganisms (Cui et al. 2022).

The significantly reduced alpha diversity of bacterial communities in diseased soils and the clear differentiation in community structure from healthy soils serve as important microbiological indicators for the occurrence of soilborne diseases in ginger. A decline in community diversity leads to reduced functional redundancy and decreased stability of microbial interaction networks, thereby substantially weakening the resistance to pathogen infection (Zhang et al. 2025). In this study, the distinct separation of microbial communities between healthy and diseased soils along the principal component analysis axis confirmed that disease occurrence is accompanied by a dramatic restructuring of the rhizosphere microbial community.

At the species composition level, beneficial genera such as *Sphingomonas* were significantly enriched in healthy soils, whereas potential pathogens such as Ralstonia dominated in diseased soils, further revealing the functional differences between healthy and diseased soils. *Sphingomonas* is a typical plant growth-promoting rhizobacterium that maintains rhizosphere health through multiple mechanisms, including siderophore secretion, solubilization of insoluble phosphates, promotion of plant growth, and antagonism against soilborne pathogens (Dor et al. 2026, Gerd et al. 2011). Previous studies have shown that this bacterial group can be recruited by plant root exudates, thereby enhancing its capacity to suppress soilborne diseases such as bacterial wilt (Zhalnina et al. 2018). *Ralstonia* is the primary pathogenic bacterium causing ginger wilt and bacterial wilt in various crops; its substantial enrichment in the rhizosphere directly indicates a decline in soil disease-suppressive capacity and a significantly increased disease risk (Su et al. 2022).

Soil pH, AN, SOM, and AP are core environmental factors driving differences in rhizosphere microbial community structure, and their regulatory roles are involved throughout the process of microecological construction. Soil pH directly exerts an environmental filtering effect on microbial taxa by altering microbial cell membrane permeability, extracellular enzyme activity, and nutrient availability (Rousk et al., 2010). Excessive AN can modify the soil carbon-to-nitrogen ratio, suppress acidophilic beneficial microbial communities, and simultaneously provide an ample nitrogen source for pathogens. As the primary carbon source for soil microorganisms, SOM is significantly positively correlated with community diversity and serves as the material foundation for maintaining the stability of beneficial microbial groups (Delgado-Baquerizo et al., 2016). Available phosphorus influences community assembly by regulating microbial energy metabolism and signal transduction pathways (Wu et al., 2025). Furthermore, the redundancy analysis and correlation analysis conducted in this study confirmed that pH and AN are positively correlated with the abundances of pathogens (*Ralstonia*), whereas SOM and AP are positively correlated with the abundances of beneficial bacteria (*Sphingomonas, Streptomyces, Ktedonobacter*), indicating that nutrient factors can regulate functional microbial communities.

In summary, soil-borne diseases of ginger result from the synergistic effects of soil nutrient imbalance and microbial community structure disturbance. This study reveals the pathogenesis of ginger from the perspective of the root-zone microenvironment and microbial interactions, providing a theoretical basis for controlling ginger soil-borne diseases through targeted regulation of soil nutrients and the rhizosphere microbial community.

## Author contributions

Shuang Ma (Conceptualization, In vestigation, Methodology, Writing original draft), Fei Fang (Soil nutrient determination), Jianming Li (Investigation), Tie Zhang and Teng Wang (Supervision, Writing-review & editing).

## Data availability

The metagenomic sequencing data generated in this study have been deposited in the NCBI Sequence Read Archive (SRA) under BioProject accession number PRJNA1455985.

